# Analysis of the Mutant Selection Window and Killing of *Mycoplasma hyopneumoniae* for Doxycycline, Tylosin, Danofloxacin, Tiamulin, and Valnemulin

**DOI:** 10.1101/704650

**Authors:** Zilong Huang, Chunxiao Mao, Yanzhe Wei, Xiaoyan Gu, Qinren Cai, Xiangguang Shen, Huanzhong Ding

## Abstract

*Mycoplasma hyopneumoniae* is the major pathogenic microorganism causing enzootic pneumonia in pigs. With increasing resistance of *M. hyopneumoniae* to conventional antibiotics, treatment is becoming complicated. Herein, we investigated the mutant selection window (MSW) of doxycycline, tylosin, danofloxacin, tiamulin, and valnemulin for treating *M. hyopneumoniae* strain (ATCC 25934) to determine the likelihood of promoting resistance with continued use of these antibiotics. Minimum inhibitory concentration (MIC) values against *M. hyopneumoniae* were determined for each antimicrobial agent and ranged from 10^5^ colony-forming units (CFU)/mL to 10^9^ CFU/mL based on microdilution broth and agar dilution methods. The minimal concentration inhibiting colony formation by 99% (MIC_99_) and the mutant prevention concentration (MPC) were determined by the agar dilution method with three inoculum sizes. Antimicrobial killing was determined based on MIC_99_ and MPC values for all five agents. MIC values ranged from 0.001 to 0.25 μg/mL based on the microdilution broth method, and from 0.008 to 1.0 μg/mL based on the agar dilution method. MPC values ranged from 0.0016 to 10.24 μg/mL. MPC/MIC_99_ values were ordered tylosin >doxycycline >danofloxacin >tiamulin >valnemulin. MPC achieved better bactericidal action than MIC_99_. Based on pharmacodynamic analyses, danofloxacin, tylosin, and doxycycline are more likely to select resistant mutants than tiamulin and valnemulin.

## 1. Introduction

*Mycoplasma hyopneumoniae* is the primary pathogen causing enzootic pneumonia, an important chronic respiratory disease in pigs resulting in high morbidity, low feed conversion rate, and considerable economic losses in the swine breeding industry **[1]**. Additionally, the disease makes pigs more susceptible to infection by secondary bacterial pathogens such as *Pasteurella multocida* and *Actinobacillus pleuropneumoniae* **[2]**. There are several kinds of antimicrobial agents that exhibit *in vitro* activity against *M. hyopneumoniae*, such as pleuromutilins, fluoroquinolones, macrolides, and tetracyclines **[3, 4]**. However, widespread use of these agents has resulted in acquired resistance of *M. hyopneumoniae* to fluoroquinolones, lincosamides, and macrolides **[5–7].** Thus, in order to reduce the risk of drug resistance, it is necessary to develop novel drugs. However, even if new drugs are discovered, re-evaluation of antimicrobial dosing is essential, and is the main method aimed at preventing the emergence and expansion of drug-resistant strains.

The mutant selection window (MSW) hypothesis postulates that, for each antimicrobial-pathogen combination, an antimicrobial concentration range exists in which selective amplification of single-step, drug-resistant mutants occurs **[8]**. The upper and lower boundaries of the MSW are the mutant prevention concentration (MPC) and the minimal concentration that inhibits colony formation by 99% (MIC_99_), respectively. The MPC is the minimum concentration that inhibits colony formation of the least antibacterial drug-susceptible mutant subpopulation. Therefore, when antimicrobial concentrations fall within the range of the MSW, this tends to lead to the enrichment of drug-resistant bacteria. Keeping drug concentrations above the MPC is likely to restrict the emergence of resistance **[9]**. This hypothesis has been verified by *in vitro* and *in vivo* experiments **[10–13].**

Because *M. hyopneumoniae* lacks a cell wall and is small bacterium (0.4 – 1.2 μm), culture isolation conditions *in vitro* are a technical challenge. In particular, quantification by the viable count method to determine colony-forming units (CFU) is arduous. Therefore, studies on the pharmacodynamics of *M. hyopneumoniae* are scarce. In the current study we determined for the first time the MPC and identified the MSW for doxycycline, tylosin, danofloxacin, tiamulin and valnemulin against *M. hyopneumoniae in vitro*. We also used time-kill tests to determine the relative antibacterial effects at the MIC_99_ and the MPC. MSW and MPC are useful parameters for optimizing dosing regimens, reducing the emergence of resistant mutants, and analyzing treatment failure **[14]**. In addition, using the CFU counting method, changes in the amount of *M. hyopneumoniae* after antibiotic action can be determined.

## 2. Materials and Methods

*Mycoplasma hyopneumoniae* standard strain ATCC 25934 was obtained as a freeze-dried powder from the China Institute of Veterinary Drug Control (Beijing, China) and stored at −80°C. Broth medium base, cysteine, and NADH were purchased from Qingdao Hope Biological Technology. Sterile swine serum was bought from Guangzhou Ruite Biological Technology. The initial pH of the medium was 7.7 ± 0.1, and 1% agar was added to solid media.

Doxycycline (85.8%), danofloxacin (100%), tiamulin (99%), tylosin (82.6%), and valnemulin (98.3%) were obtained from Guangdong Dahuanong Animal Health Products. These five antibacterial agents were dissolved in Milli-Q water and sterilized by filtration. A 1280 μg/mL fresh stock solution of each antibacterial agent was prepared for each experiment.

### 2.1 Determination of minimum inhibitory concentration (MIC)

MIC values were determined as described previously **[15]**. Briefly, MIC values were calculated for 10^5^, 10^6^, and 10^7^ CFU/mL *M. hyopneumoniae* cultures in the exponential phase. A 100 μL sample of exponential phase culture was added to an equal volume of drug-containing medium culture in a 96-well plate. A growth control (inoculum without antimicrobials), sterility control (sterile broth at pH 7.8), and end-point control (blank medium at pH 6.8) were included. Plates were cultured at 37°C with 5% CO_2_ in an incubator after being sealed. When the color of the growth control was the same as the end-point control, the MIC was determined as the minimal concentration of antibacterial agent that resulted in no color change.

MIC values were determined by the agar dilution method as described previously **[16]**. A 10 μL sample of *M. hyopneumoniae* culture (10^5^–10^7^ CFU) was placed on the surface of a plate in which wells contained 1.25–20 μg/mL danofloxacin, 2–32 μg/mL tiamulin, 4–64 μg/mL tylosin, 5–80 μg/mL doxycycline, or 0.16–2.56 μg/mL for two-fold agar dilution analysis. All plates were incubated for at least 8 days. Meanwhile, growth control plates without antimicrobials were set up for each test, and all experiments were repeated three times. The lowest concentration without *M. hyopneumoniae* growth on agar plates was taken as the MIC value. Each test was repeated three times.

### 2.2 Measurement of MIC_99_ and mutant prevention concentration (MPC)

MIC_99_ values were measured as reported previously **[17]** with modifications. MIC_99_ drug concentrations were based on linear decreasing dilutions of MIC values. The antibiotic concentration ranged from 1 × MIC to 0.5 × MIC in sequential 10% dilution decreases. The quantity of bacteria in the logarithmic growth phase reached 10^7^ CFU/mL. Three 10 μL drops of each diluted suspension were inoculated onto agar plates and cultured for at least 8 days as described above. Colony numbers between 30 and 300 were counted.

The MPC is defined as the lowest drug concentration that prevents bacterial colony formation from a culture containing ≥10^9^ CFU/mL bacteria **[18].** We attempted different centrifugal methods for enriching *M. hyopneumoniae*. Ultimately, an 800 mL stationary growth phase culture was transferred into 20 tubes (each 50 mL), tubes were centrifuged (5000 × g for 20 min), and each bacteria solution was resuspended in 1 mL fresh medium. All 20 enriched cultures were combined into two 15 mL tubes, centrifuged (5000 × g for 20 min), and resuspended in 1 mL fresh medium for counting. The final concentration of *M. hyopneumoniae* was 8.8×10^9^ CFU/mL.

MPC values were measured by the agar method as described previously **[19].** Briefly, 200 μL samples of each enriched culture were inoculated onto agar plates containing various concentrations of antibiotic (six parallel solid plates per antibiotic concentration). These plates were incubated at 37°C with 5% CO_2_ in a humidified incubator for 8–10 days. The lowest antibiotic concentration that resulted in no colony formation was considered the primary MPC (MPC_pr_). After a 20% linear drug concentration decrease in MPC_pr_, the MPC was tested again, and recorded as the lowest drug concentration preventing bacterial growth. Each test was repeated three times.

### 2.3 Time-kill tests

*In vitro* time-killing assays were performed as described previously **[20]**. Briefly, MIC_99_ and MPC values for all five agents were tested. After adding 3.5 mL blank medium and 0.1 mL drug solution (40 times the target concentration) to each penicillin bottle, 0.4 mL exponential *M. hyopneumoniae* suspension with an inoculum size between 10^5^ CFU/mL and 10^9^ CFU/mL was added. Cultures were incubated at 37°C with 5% CO_2_ for 48 h. Aliquots of 100 μL were collected from each culture at 0, 3, 6, 9, 12, 24, 36, and 48 h. The viable cell number was determined via 10-fold serial dilutions and plating 10 μL of each diluted culture on drug-free agar. Growth controls (*M. hyopneumoniae* cultures without drugs) and sterility controls (5 mL medium at pH 7.8) were also included. Plates were incubated for at least 8 days at 37°C with 5% CO_2_ in a humidified incubator. Each test was repeated three times.

## 3. Results

### 3.1 MIC determination

MICs of danofloxacin, tiamulin, tylosin, doxycycline, and valnemulin against *M. hyopneumoniae* determined by the microdilution and agar dilution methods are shown in **Figure 1**. Values determined using the solid MIC method were 8-fold higher (tylosin), 4-fold higher (danofloxacin, doxycycline, and valnemulin), and 2-fold higher (tiamulin) than those determined by the liquid method at an identical inoculum size of 10^5^ CFU/mL. In addition, as the inoculum size used in these assays was increased, the MIC values also increased. *M. hyopneumoniae* displayed its greatest sensitivity to valnemulin and its least to doxycycline

**Figure 1.**
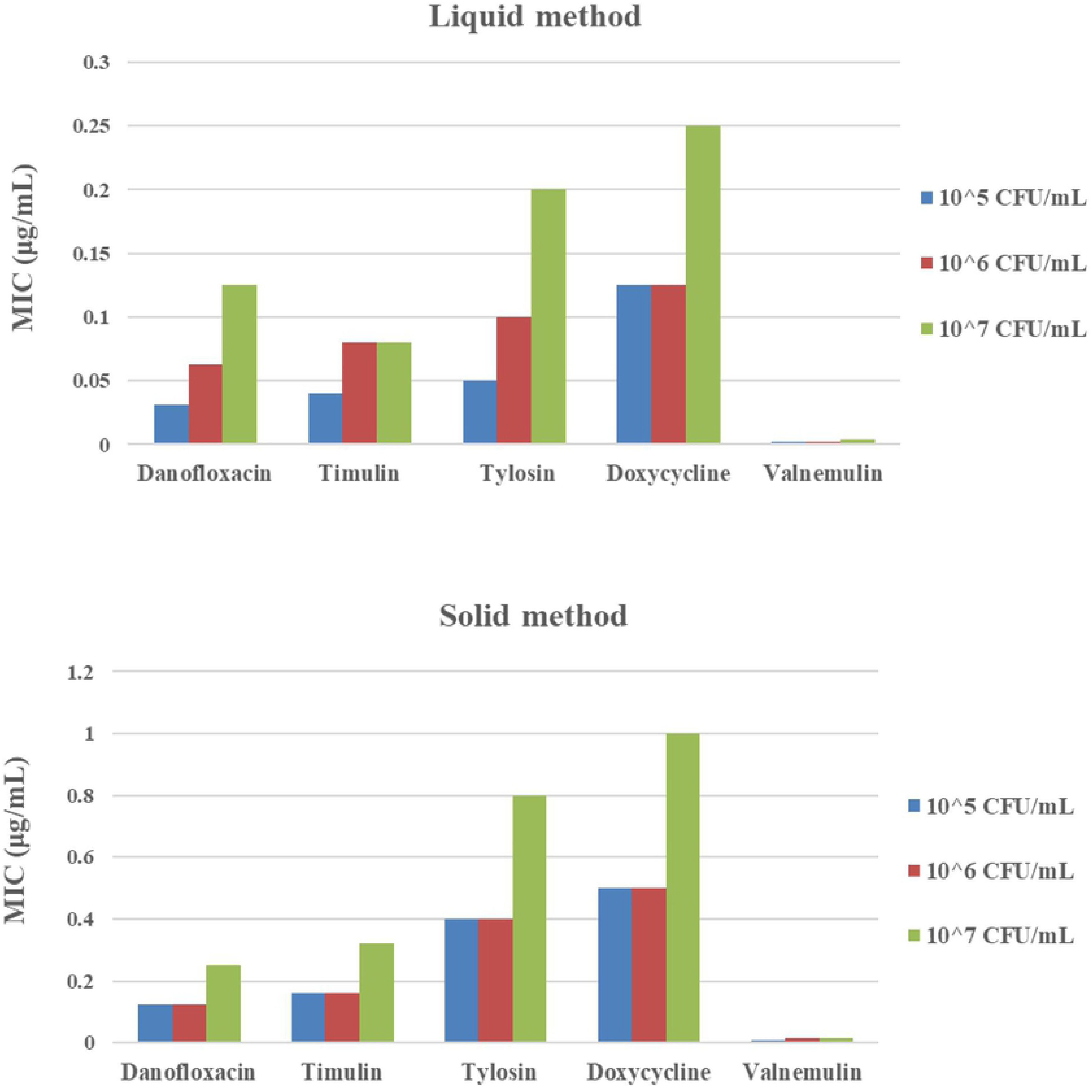
Minimum inhibitory concentration (MIC) determination for danofloxacin, tiamulin, tylosin, doxycycline, and valnemulin against *M. hyopneumoniae* in artificial medium using liquid and solid agar methods with inoculum sizes of 10^5^, 10^6^, and 10^7^ CFU/mL.

### 3.2 Determination of MIC_99_, MPC, and selection index (SI)

MIC_99_, MPC, and SI values are shown in **Table 1**. The SI, the ratio of MPC to MIC_99_, reflects the ability of an antibacterial agent to induce resistant mutants. MIC_99_ values for all antibacterial agents ranged from 0.0122 to 0.343 μg/mL, and MPC values ranged from 0.016 to 10.24 μg/mL. SI values ranged from 1.31 to 10.24, and were ranked valnemulin <tiamulin <danofloxacin <doxycycline <tylosin (low to high). Thus, valnemulin displayed the lowest SI value, while tylosin exhibited the highest SI value.

**Table 1.**
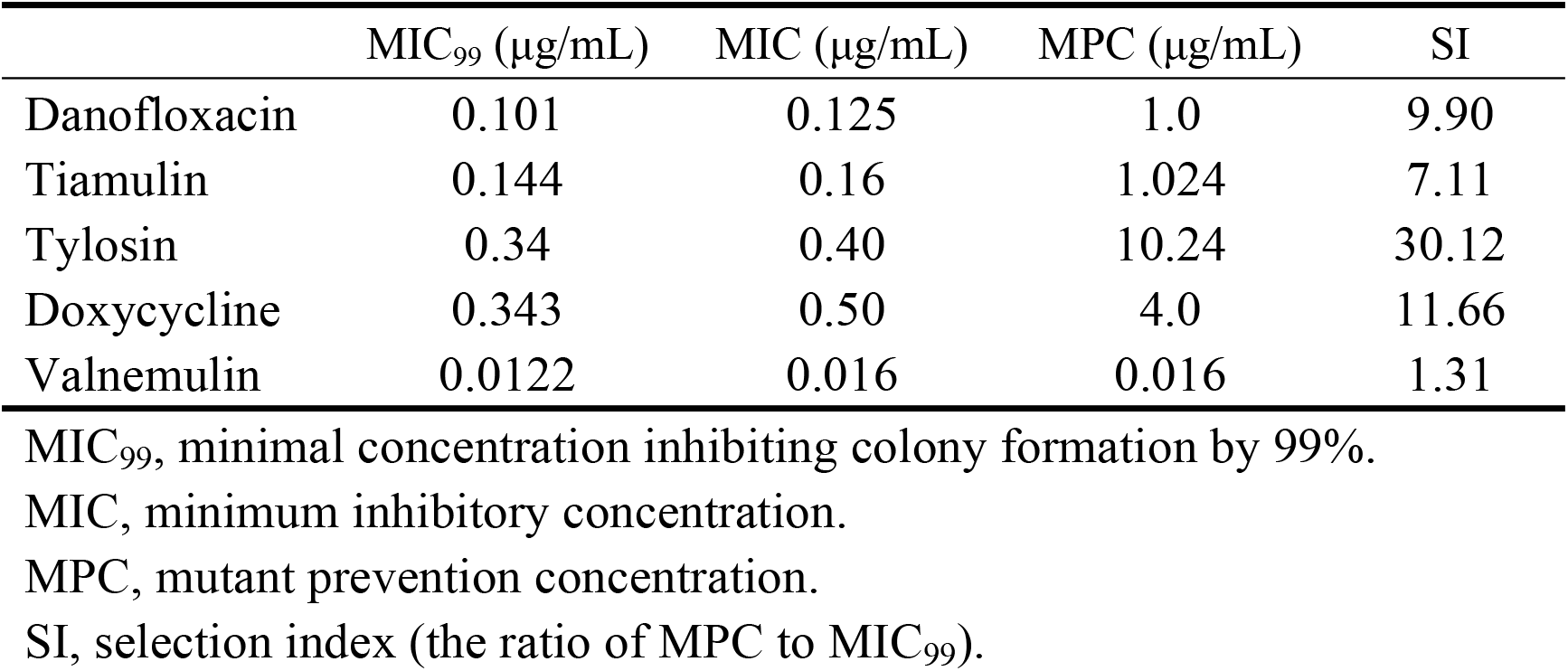
Comparison of MIC_99_, MIC, MPC, and SI values for five antimicrobial agents tested against *M. hyopneumoniae*

### 3.3 In vitro killing analysis

Time-kill curves of compounds against *M. hyopneumoniae* were obtained using three different inoculum sizes. Reductions in *M. hyopneumoniae* count with different inoculum sizes for MIC_99_ and MPC are listed in **Table 2 and Table 3**. Danofloxacin, tiamulin, tylosin, and valnemulin achieved bactericidal activity against 10^5^ CFU/mL *M. hyopneumoniae* with MIC_99_ dosage, while doxycycline achieved bacteriostatic activity only. Colony count reductions recorded at the 48h time point 3.63, 3.68, 3.75, and 3.61 log_10_ CFU/mL for danofloxacin, tiamulin, tylosin, and valnemulin, respectively, but only 1.4 log_10_ CFU/mL for doxycycline. All five compounds achieved bactericidal activity against 10^5^ CFU/mL *M. hyopneumoniae* with MPC dosage **(Figure 2)**.

**Table 2.**
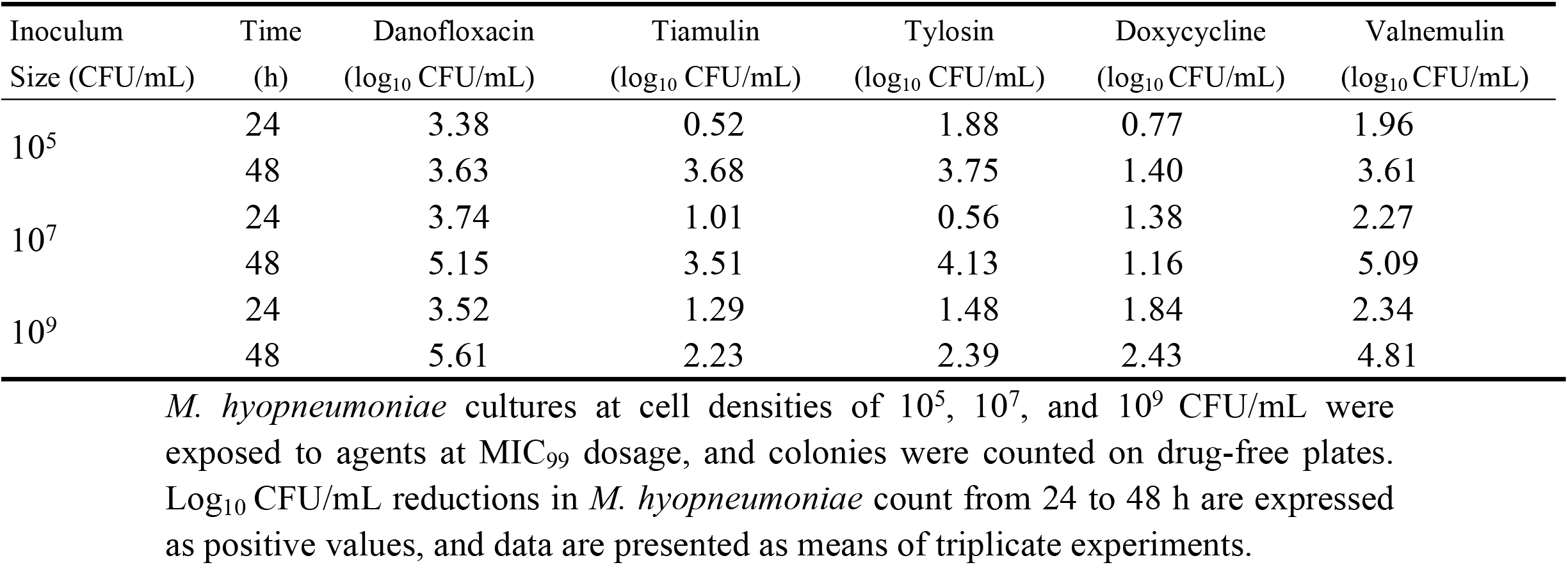
Reduction in *M. hyopneumoniae* growth for three different inoculum sizes based on measured MIC_99_ concentrations

**Table 3.** Reduction in *M. hyopneumoniae* growth for three different inoculum sizes based on measured MPC concentrations

| Inoculum size (CFU/mL) | Time (h) | Danofloxacin (log<sub>10</sub> CFU/mL) | Tiamulin (log<sub>10</sub> CFU/mL) | Tylosin (log<sub>10</sub> CFU/mL) | Doxycycline (log<sub>10</sub> CFU/mL) | Valnemulin (log<sub>10</sub> CFU/mL) |
| --- | --- | --- | --- | --- | --- | --- |
| 10<sup>5</sup> | 24 | 3.63 | 2.25 | 2.65 | 2.83 | 3.43 |
|  | 48 | 3.63 | 3.68 | 3.75 | 3.72 | 3.61 |
| 10<sup>7</sup> | 24 | 3.08 | 3.16 | 2.47 | 2.57 | 2.99 |
|  | 48 | 5.15 | 5.21 | 5.13 | 4.16 | 5.09 |
| 10<sup>9</sup> | 24 | 4.63 | 3.48 | 2.43 | 2.62 | 3.47 |
|  | 48 | 7.03 | 4.13 | 4.36 | 3.96 | 4.90 |

*M. hyopneumoniae* cultures at cell densities of 10<sup>5</sup>, 10<sup>7</sup>, and 10<sup>9</sup> CFU/mL were exposed to agents at MIC<sub>99</sub> dosage, and colonies were counted on drug-free plates. Log<sub>10</sub> CFU/mL reductions in *M. hyopneumoniae* count from 24 to 48 h are expressed as positive values, and data are presented as means of triplicate experiments.

**Figure 2.**
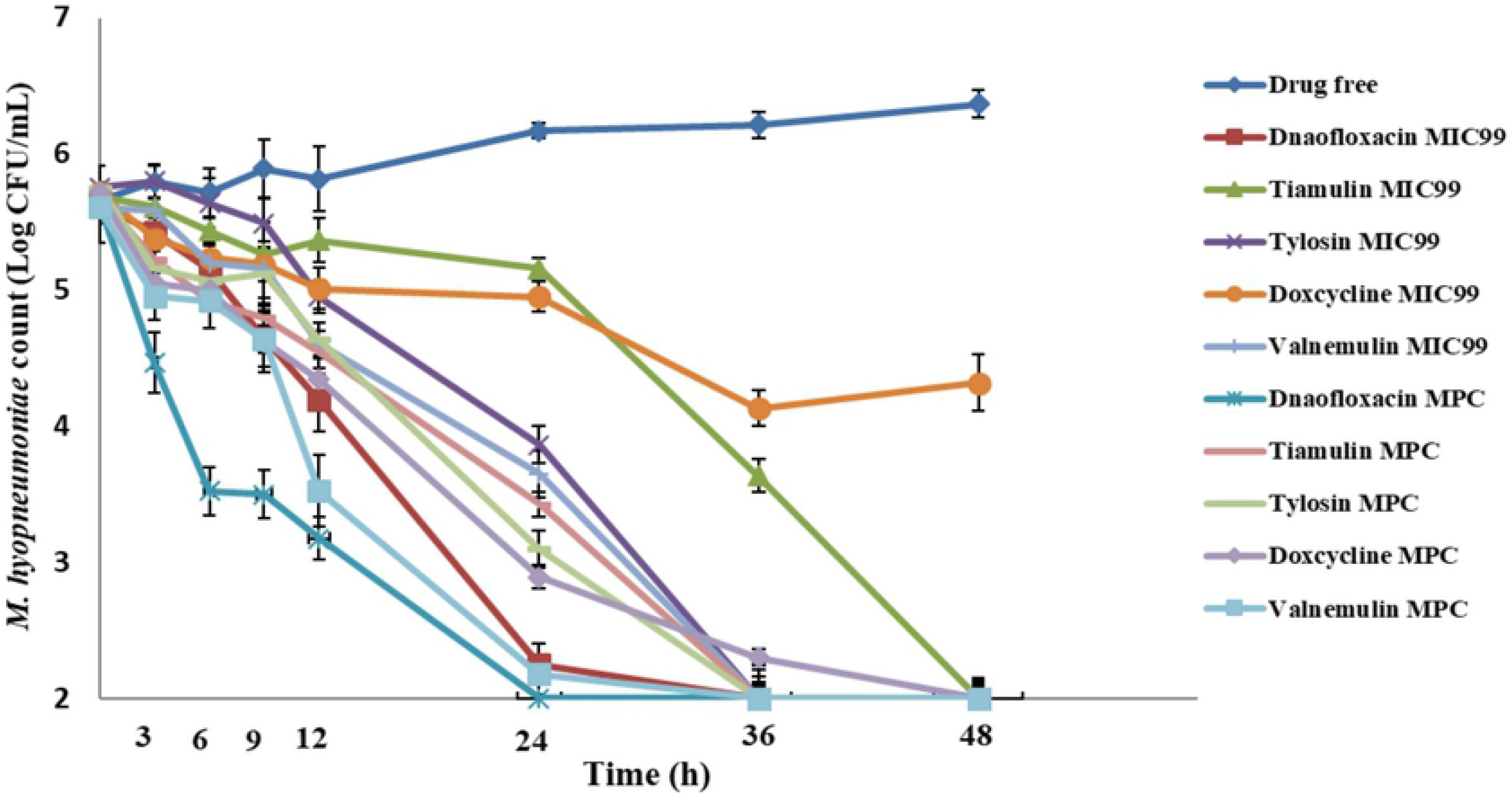
*M. hyopneumoniae* killing curves at the minimal concentration inhibiting colony formation by 99% (MIC_99_) and at the mutant prevention concentration (MPC) with an inoculum of 10^5^ CFU/mL.

At a greater inoculum size (10^7^ CFU/mL), danofloxacin, tiamulin, tylosin and valnemulin were bactericidal at the MIC_99_ while doxycycline was bacteriostatic only. Colony count reductions recorded at the 48h time point were 5.15 log_10_ CFU/mL for danofloxacin, 5.09 log_10_ CFU/mL for valnemulin, 3.51 log_10_ CFU/mL for tiamulin, 4.13 log_10_ CFU/mL for tylosin, and 1.16 log_10_ CFU/mL for doxycycline. All five compounds achieved bactericidal activity against 10^7^ CFU/mL *M. hyopneumoniae* with MPC dosage **(Figure 3)**.

**Figure 3.**
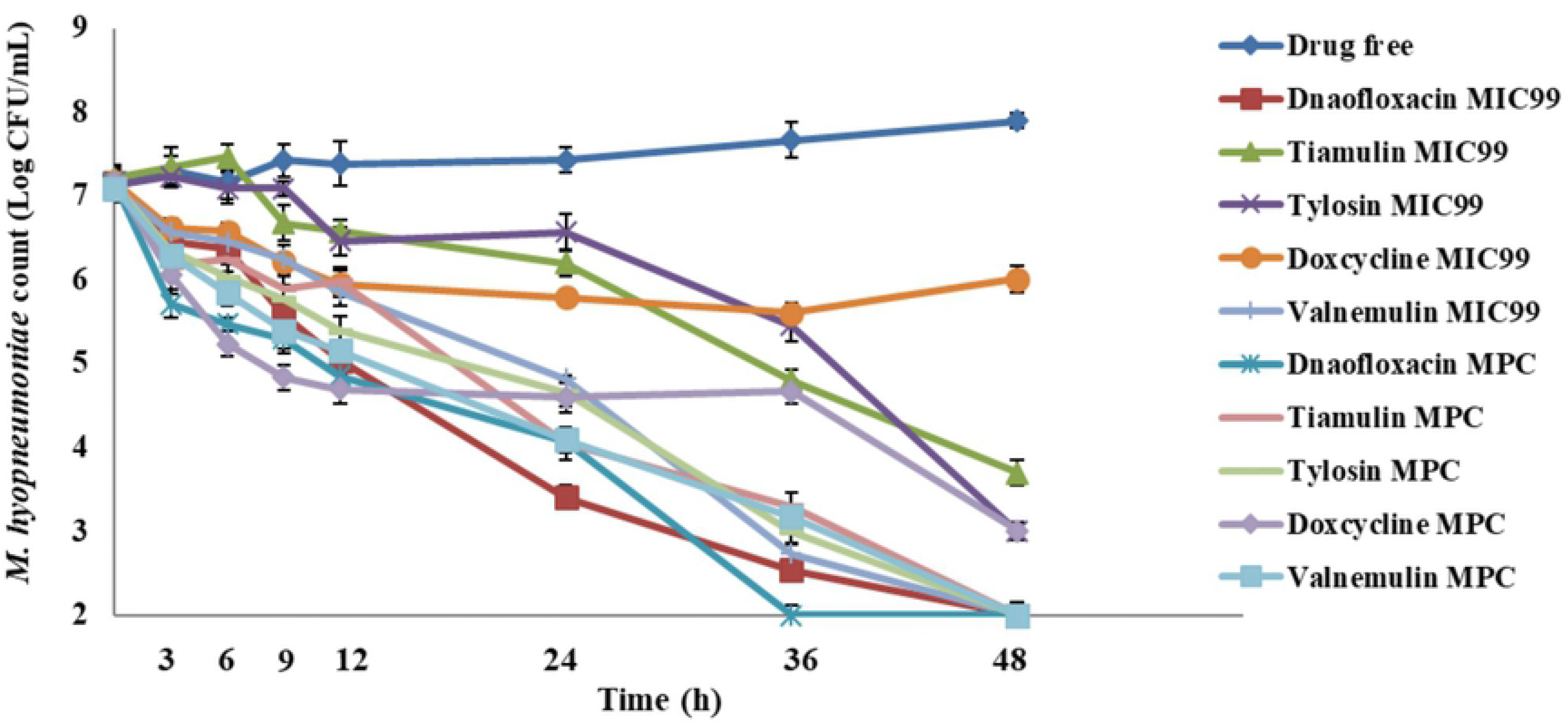
*M. hyopneumoniae* killing curves at the minimal concentration inhibiting colony formation by 99% (MIC_99_) and at the mutant prevention concentration (MPC) with an inoculum of 10^7^ CFU/mL.

Danofloxacin and valnemulin both achieved bactericidal activity against 10^9^ CFU/mL of *M. hyopneumoniae* cells at the MIC_99_ concentrations while the other three were bacteriostatic only. Colony count reductions recorded at 48 h were danofloxacin 5.61 log_10_ CFU/mL, valnemulin 4.81 log_10_ CFU/mL, tiamulin 2.23 log_10_ CFU/mL, tylosin 2.39 log_10_ CFU/mL and doxycycline 2.43 log_10_ CFU/mL. All compounds achieved bactericidal activity against 10^9^ CFU/mL of *M. hyopneumoniae* at the MPC concentrations. Overall, the rank order of antibacterial agents for colony count reduction was danofloxacin >valnemulin >tylosin >tiamulin >doxycycline **(Figure 4)**.

**Figure 4.**
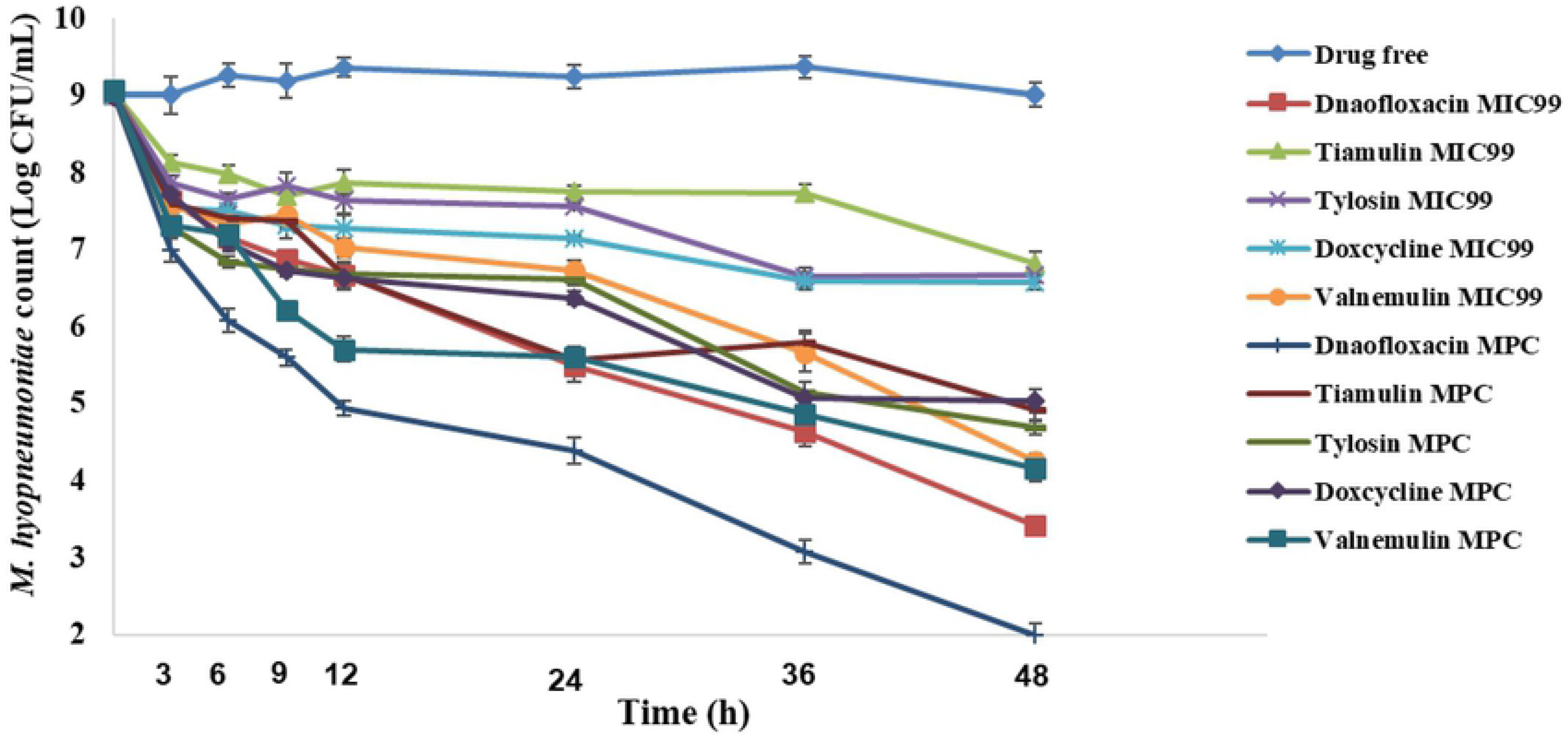
*M. hyopneumoniae* killing curves at the minimal concentration inhibiting colony formation by 99% (MIC_99_) and at the mutant prevention concentration (MPC) with an inoculum of 10^9^ CFU/mL.

## 4. Discussion

*M. hyopneumoniae* is a major respiratory disease-causing pathogen in modern intensive pig farming worldwide. Although vaccine-based immunization is an important preventive measure for enzootic pneumonia, treatment with antibacterial agents is known to accelerate disease recovery and reduce disease-related complications. However, strains resistant to enrofloxacin and tylosin have appeared among clinically isolated strains **[6, 7]**. In particular, many fluoroquinolones are important antibiotics for the treatment of human infections, and are more likely to lead to cross-resistance. Due to the difficulties associated with *in vitro* culturing and viability counting (CFU measurements) for *M. hyopneumoniae*, systematic *in vitro* pharmacodynamic evaluation of antibacterial agents against *M. hyopneumoniae* is scarce. Thus, in the present study, *in vitro* pharmacodynamic indices of several representative antimicrobial agents against *M. hyopneumoniae* were determined, and killing curves were plotted. To the best of our knowledge, this is the first study to explore the risk of *M. hyopneumoniae* resistance using MPC and MSW parameters.

MIC values determined by the liquid method were similar to those measured in previous studies **[4, 5]**. Using both liquid and solid agar methods, MIC values increased with increasing inoculum size; values obtained with a large inoculum were two to four times higher than those obtained with small inoculum. It has been reported previously that MIC values increase with increasing bacterial load **[21, 22]**. Among the five antibiotics tested, danofloxacin and tylosin were more sensitive to inoculum size for MIC determination. In an earlier report **[23],** a larger inoculum of *Staphylococcus aureus* had a more significant effect on the antibacterial activity of nafcillin and vancomycin than a smaller inoculum. Therefore, in cases of high bacterial inoculation, more careful consideration is required when selecting the MIC reference value for the relevant experiment. In clinical treatment, the MIC value should be determined for different bacterial counts according to the severity of animal infection to establish a better treatment plan.

The MPC is defined as the concentration of antibacterial drug that prevents the growth of large quantities of resistant sub-populations. Because culturing of *M. hyopneumoniae* is difficult *in vitro*, determination of MPC values is challenging; after much effort, *M. hyopneumoniae* was cultured to a cell density of 10^7^–10^8^ CFU/mL. We tried a variety of enrichment methods to generate quantities of bacteria sufficient for determining MPCs, and eventually managed to measure MPCs for all five antibiotics. Relationships between antimicrobial exposure, MPC/MSW values and antimicrobial resistance selection have been explored previously. For example, in *Staphylococcus aureus*, cefquinome concentrations below the MIC_99_, intermediate between the MIC_99_ and MPC, and above the MPC resulted in the selection of mutants that differed in terms of the proportion of resistant and susceptible bacteria **[11]**. When the concentration of levofloxacin fell within the MSW, the sensitivity of *S. aureus* decreased and mutant subpopulations emerged **[24]**. For *M. hyopneumoniae*, the reported danofloxacin, tylosin, and doxycycline concentrations in pigs fell within the MSW completely after an intramuscular dose of 2.5 or 10 mg/kg (body weight); after oral administration at a dose of 20 mg/kg, C_max_ values for danofloxacin, tylosin, and doxycycline were 0.45 ± 0.09, 2.71 ± 1.09, and 2.44 ± 0.51 μg/mL **[25–27]**. Correlative MIC_99_ and MPC values were 0.101 and 1.0, 0.34 and 10.24, and 0.343 and 4.0 μg/mL, respectively. The reported tiamulin and valnemulin concentrations were comfortably above the MSW values only after oral administration at a dose of 10 or 25 mg/kg **[28, 29]**. Consequently, by combining the *in vivo* pharmacokinetic parameters and the *in vitro* pharmacodynamic results, we speculate that danofloxacin, tylosin, and doxycycline are more likely to select resistant mutants than tiamulin and valnemulin. Continued use of first-line antibacterial agents against *M. hyopneumoniae* according to current dosing regimens may therefore promote drug resistance selection, and hence limit their long-term efficacy in the treatment of endemic pneumonia in pigs.

We determined the bactericidal effects of danofloxacin, tylosin, doxycycline, tiamulin, and valnemulin against *M. hyopneumoniae* at various bacterial densities and drug concentrations. At three different inoculation amounts, doxycycline displayed bacteriostatic activity at MIC_99_ dosage and bactericidal action at MPC dosage. At the highest inoculation amount, tiamulin and valnemulin acted as bacteriostatic agents at MIC_99_ dosage and as bactericidal agents at MPC dosage. Danofloxacin exhibited the fastest sterilization rate. Moreover, valnemulin was highly sensitive to *M. hyopneumoniae*, and exerted an obvious bactericidal effect. These results showed that when the concentration of antibiotic equaled or exceeded the MPC, *M. hyopneumoniae* was rapidly killed. Drug concentrations at the MPC also reduced the chances of bacteria re-growing during drug exposure. These results are similar to those of an earlier report **[30]**. In this previous study, the bactericidal effect at MIC was slow and incomplete. However, at MPC and maximum serum or tissue drug concentrations, killing was more pronounced than at MIC, and increased with increasing duration of drug exposure.

The main limitation of the present study was that the *in vitro* pharmacodynamic determination of *M. hyopneumoniae* was only carried for the standard strain, and clinical isolates should be assessed to confirm our findings. Nevertheless, the present work represents a meaningful pilot study in this area. A second limitation is that all experiments were performed under ideal conditions *in vitro*, without considering the complexity of factors *in vivo*. Thus, *in vivo* experiments are currently being explored.

In conclusion, the present study was the first to establish pharmacodynamic analyses of five antimicrobial agents against *M. hyopneumoniae*. And we determined MPC and MSW parameters to explore the risk of *M. hyopneumoniae* resistance. The results showed that the bactericidal action of MPC was better than MIC_99_, and the antibacterial effects of these drugs against *M. hyopneumoniae* are significantly different. These pharmacodynamic results are meaningful in choosing antimicrobials for therapy. And danofloxacin, tylosin, and doxycycline are more likely to select resistant mutants than tiamulin and valnemulin.

## Acknowledgments

This work was supported by the National Key Research and Development Program of China (grant numbers 2016YFD0501300 and 2016YFD0501310).

## Author Contributions

Methodology, software, validation, formal analysis, data curation, manuscript preparation, manuscript reviewing and editing, visualization, and project administration were performed by ZH. ZH, CM, ZZ, and LZ contributed to investigations. Resources were provided by XG, QC, XS, and HD. Supervision was provided by HD, and funding was acquired by HD.

